# The tumor microbiome reacts to hypoxia and can influence response to radiation treatment in colorectal cancer

**DOI:** 10.1101/2023.09.01.555936

**Authors:** Martin Benej, Rebecca Hoyd, McKenzie Kreamer, Caroline E. Wheeler, Dennis J. Grencewicz, Fouad Choueiry, Carlos H.F. Chan, Yousef Zakharia, Qin Ma, Rebecca D. Dodd, Cornelia M. Ulrich, Sheetal Hardikar, Michelle L. Churchman, Ahmad A. Tarhini, Lary A. Robinson, Eric A. Singer, Alexandra P. Ikeguchi, Martin D. McCarter, Gabriel Tinoco, Marium Husain, Ning Jin, Aik Choon Tan, Afaf E.G. Osman, Islam Eljilany, Gregory Riedlinger, Bryan P. Schneider, Katarina Benejova, Martin Kery, Ioanna Papandreou, Jiangjiang Zhu, Nicholas Denko, Daniel Spakowicz

## Abstract

Tumor hypoxia has been shown to predict poor patient outcomes in several cancer types, partially because it reduces radiation’s ability to kill cells. We investigated whether some of the clinical effects of hypoxia could also be due to its impact on the tumor microbiome. We examined the RNA-seq data from the Oncology Research Information Exchange Network (ORIEN) database of colorectal cancer (CRC) patients treated with radiotherapy. For each tumor, we identified microbial RNAs and related them to the hypoxic gene expression scores calculated from host mRNA. Our analysis showed that the hypoxia expression score predicted poor patient outcomes and identified tumors enriched with certain microbes such as *Fusobacterium nucleatum*. The presence of other microbes, such as *Fusobacterium canifelinum,* predicted poor patient outcomes, suggesting a potential interaction between hypoxia, the microbiome, and radiation response. To investigate this concept experimentally, we implanted CT26 CRC cells into both immune-competent BALB/c and immune-deficient athymic nude mice. After growth, where tumors passively acquired microbes from the gastrointestinal tract, we harvested tumors, extracted nucleic acids, and sequenced host and microbial RNAs. We stratified tumors based on their hypoxia score and performed metatranscriptomic analysis of microbial gene expression. In addition to hypoxia-trophic and -phobic microbial populations, analysis of microbial gene expression at the strain level showed expression differences based on the hypoxia score. Hypoxia appears to not only associate with different microbial populations but also elicit an adaptive transcriptional response in intratumoral microbes.

**SIGNIFICANCE:** Tumor hypoxia reduces radiation’s ability to kill cells. We explored whether some of the clinical effects of hypoxia could also be due to interaction with the tumor microbiome. Hypoxic expression scores associated with certain microbes and elicited an adaptive transcriptional response in others.

## INTRODUCTION

Colorectal cancer (CRC) is the third most common and second most lethal cancer worldwide, accounting for 1.9 million cases and nearly 900,000 deaths every year (1). More than 60% of all CRC cases occur in the sigmoid colon and are diagnosed at stage II or above (2). The 5-year survival rate for CRC typically ranges from 90% for patients diagnosed with localized disease to 14% for those diagnosed with metastatic disease (3). Clinical management of CRC depends on the stage: surgery is the main treatment option in the early stages, while neoadjuvant chemoradiation therapy (nCRT) followed by surgery is the standard of care for locally advanced stages (4). However, only about 15% of CRC patients treated with nCRT achieve complete pathological responses (5). Because dose escalation dramatically increases the risk of toxicity and exceeds the radiotolerance of adjacent normal tissues, understanding the mechanisms behind CRC resistance to nCRT therapy is of utmost clinical importance.

The human intestinal microbiome comprises 10^13^-10^14^ mutualistic microorganisms that play a crucial role in shaping the intestinal epithelium, harvesting nutrients, maturation of the host immune system, defense against pathogens, and maintenance of gut barrier function (6–10). Increasing lines of evidence suggest that dysbiosis or alteration of the intestinal microbiome composition and function is increasingly involved with the initiation and progression of CRC (11). Moreover, emerging studies suggest a direct association between intestinal microbiome dysbiosis and sensitivity to anti-cancer therapy (12–15). In particular, studies in gnotobiotic mice have shown that intestinal microbiota can shape the response to anti-cancer therapy, suggesting complex host-microbiome crosstalk (16,17).

The microenvironment of the human gastrointestinal (GI) tract is generally hypoxic (physiological hypoxia), showing steep oxygen gradients along the radial axis ranging from well-oxygenated subepithelial mucosa to the anaerobic lumen (18,19). The lumen is inhabited by a vast number of anaerobic microbes that thrive under anoxic conditions. Microbial colonization of CRC tumors by microbial taxa is therefore highly influenced by the route by which they reach the tumor and by the ability of the microorganism to survive the environmental oxygen within the tumor microenvironment (TME). Because hypoxia directly limits the efficiency of radiation therapy (20), we have asked whether potential crosstalk between the CRC tumor microbiome and the hypoxic TME can influence the tumor response to radiotherapy.

In this study, we analyzed RNA-seq data from 141 pretreatment CRC samples from patients receiving radiation therapy as a part of their treatment regimen to identify environmental hypoxia- and radiation treatment-dependent variations in the tumor microbiome. We then assessed these patterns for their impact on overall survival. We performed in vivo validation of these concepts in heterotopic model CT26 tumors to identify microbiome composition and gene expression as a function of tumor oxygenation. Comparison of host animals from BALB/c and athymic nude strains identified hypoxia-dependent microbial populations and adaptive gene-level response to varying oxygen conditions.

## METHODS

### Study Design

The Oncology Research Information Exchange Network (ORIEN) is an alliance of 18 US cancer centers established in 2014. All ORIEN alliance members utilize a standard Total Cancer Care® (TCC) protocol. As part of the TCC study, participants agree to have their clinical data followed over time, to undergo germline and tumor sequencing, and to be contacted in the future by their provider if an appropriate clinical trial or other study becomes available (21). TCC is a prospective cohort study with a subset of patients enrolled in the ORIEN Avatar program, which includes research-use-only (RUO)-grade whole-exome tumor sequencing, RNA sequencing, germline sequencing, and collection of deep longitudinal clinical data with lifetime follow-up. Nationally, over 325,000 participants have enrolled in TCC. M2GEN, the commercial and operational partner of ORIEN, harmonizes all abstracted clinical data elements and molecular sequencing files into a standardized, structured format to enable aggregation of de-identified data for sharing across the network. This study included 2,755 ORIEN Avatar patients diagnosed with melanoma, sarcoma, thyroid, pancreatic, colorectal, or lung cancer who consented to the TCC protocol from the participating members of ORIEN. Of these, 500 patients had colon adenocarcinoma and 95 had rectal adenocarcinoma.

### Sequencing Methods

ORIEN Avatar specimens undergo nucleic acid extraction and sequencing at HudsonAlpha (Huntsville, AL) or Fulgent Genetics (Temple City, CA). For frozen and optimal cutting temperature (OCT) tissue RNA extraction, Qiagen RNAeasy plus mini kit is performed, generating 216 bp average insert size. For formalin-fixed paraffin-embedded (FFPE) tissue, Covaris Ultrasonication FFPE DNA/RNA kit is utilized to extract both DNA and RNA, generating a 165 bp average insert size. RNA sequencing (RNA-seq) is performed using the Illumina TruSeq RNA Exome with single library hybridization, cDNA synthesis, library preparation, and sequencing (100 bp paired reads at Hudson Alpha, 150 bp paired reads at Fulgent) to a coverage of 100M total reads/50M paired reads. RNA-seq tumor pipeline analysis is processed according to the workflow outlined below using GRCh38/hg38 human genome reference sequencing and GenCode build version 32. Adapter sequences are trimmed from the raw tumor sequencing FASTQ file. Adapter trimming via k-mer matching is performed along with quality trimming and filtering, contaminant filtering, sequence masking, guanine-cytosine (GC) filtering, length filtering, and entropy filtering. The trimmed FASTQ file is used as input to the read alignment process. The tumor adapter-trimmed FASTQ file is aligned to the human genome reference (GRCh38/hg38) and the Gencode genome annotation v32 using the STAR aligner. The STAR aligner generates multiple output files used for gene fusion prediction and gene expression analysis. RNA expression values are calculated and reported using estimated mapped reads, fragments per kilobase of transcript per million mapped reads (FPKM), and transcripts per million mapped reads (TPM) at both the transcript and gene levels based on transcriptome alignment generated by STAR. For model murine tumors, we used Powerfecal pro kits from Qiagen to extract RNA and DNA.

### Athymic Nude and BALB/c Mice Tumors

Animals were purchased from Charles River (housed in groups of 5) and were injected subcutaneously on the right flank with 0.5 million CT26 cells with tumor growth measured by calipers using the formula (L×W^2^)/2. Radiotherapy was delivered by a single tangential beam from the Small Animal Radiation Research Platform (SARRP). Untreated tumors were typically harvested on day 16 after sacrifice when most tumors were at removal criteria. Mice stool was collected before tumor harvesting.

### Hypoxia Score Generation

Hypoxia scores were generated using the R package (22) for Buffa, Winter, and Leonard signatures. The R package {mt.surv} was used to compare these scores’ relationships to overall survival, and Buffa (23) was chosen to represent hypoxia in further analyses. For all analyses comparing low and high hypoxia scores, samples were defined as having low hypoxia if their hypoxia score was in the lower tertile of the data and high hypoxia if their score was in the upper tertile of the data.

Kaplan-Meier (KM) survival curves were generated, and Cox proportional hazards models were applied using the R package {survival}.

### Differential Abundance and Expression Analyses

The R package {DESeq2} was used to compare the abundance of microbes in low- and high-hypoxia tumors for both The Cancer Genome Atlas (TCGA) and ORIEN data. It was also used to compare microbe abundances in low- and high-hypoxia tumors in the mouse experiments, as well as the behavior of microbe gene expressions in high- and low-hypoxia mouse tumors.

### Mouse Sequencing Processing – Microbe Genes

To obtain information on the gene expression of microbes in the mouse experiments, RNA-seq data from the mouse tumors were classified using HUMAnN 3.0 with MetaPhlAn and the default ChocoPhlAn database.

### {exotic} Processing

Microbes were classified in both human and mouse samples using the {exotic} pipeline. This entails first removing as many human or mouse reads as possible by aligning to the appropriate genome using STAR. The hg38 genome was used for human samples, the GRC m39 genome was used for mouse samples, and annotated genes were further used in gene expression analyses. Reads left unclassified after STAR alignment were then aligned to a Kraken 2 database customized to include fungi and archaea, after which Bracken (short for Bayesian reestimation of abundance with Kraken) was used to determine the likely species of all classified reads. The R package {exotic} (24) was then used to decontaminate the results and to normalize the human samples to account for processing at different sequencing centers.

### Microbial Culture and Healthy Gut Mixture

Representative bacteria strains *Akkermansia municiphila* (ATCC^®^ BAA-35™), *Lactobacillus acidophila* (ATCC^®^ 4356™), *Streptococcus thermophilus* (ATCC^®^ 19258™), *Bacteroides ovatus* (ATCC^®^ 8483™), and *Dorea formicigenerans* (ATCC^®^ 27755™) were obtained from American Type Culture Collection (ATCC, Manassas, VA). Fecal bacteria isolates were collected from 3 healthy adult volunteers as described previously (25) (protocol approval was obtained from the Institutional Review Board, and informed consent was documented for each individual). All bacterial strains were maintained in standard culture conditions as previously described (26). Briefly, overnight cultures of bacteria were established in Gifu anaerobic medium (GAM) (HiMedia, West Chester, PA). The cultures were maintained in an anaerobic environment at 37°C using a Coy Laboratories anaerobic chamber (Coy Lab, Flint, MI). The final optical density (OD) of bacterial cultures was measured using an ELx808 absorbance plate reader (BioTek, Winooski, VT) to estimate growth. Drop plate analysis was also used to quantify the colony-forming units (CFU) in each culture. Prior to injections, bacteria cultures were centrifuged and then resuspended at a final concentration of 3×10^9^ CFU per injection.

## RESULTS

Virtually all solid tumor types are characterized by some level of hypoxia. One way to approximate tumor hypoxia is to use quantitative gene expression signatures of hypoxia-regulated genes. For **Figure 1A**, we calculated the values and distribution of the hypoxia expression score (23) across cancer types within the ORIEN dataset (23). This hypoxia signature is similar to others and scores sets of genes induced in response to hypoxia primarily by the HIF1 transcription factor. This gene family can be extracted from tumor mRNA data in publicly available clinical datasets to identify tumors with significant levels of hypoxia.

**Figure 1.**
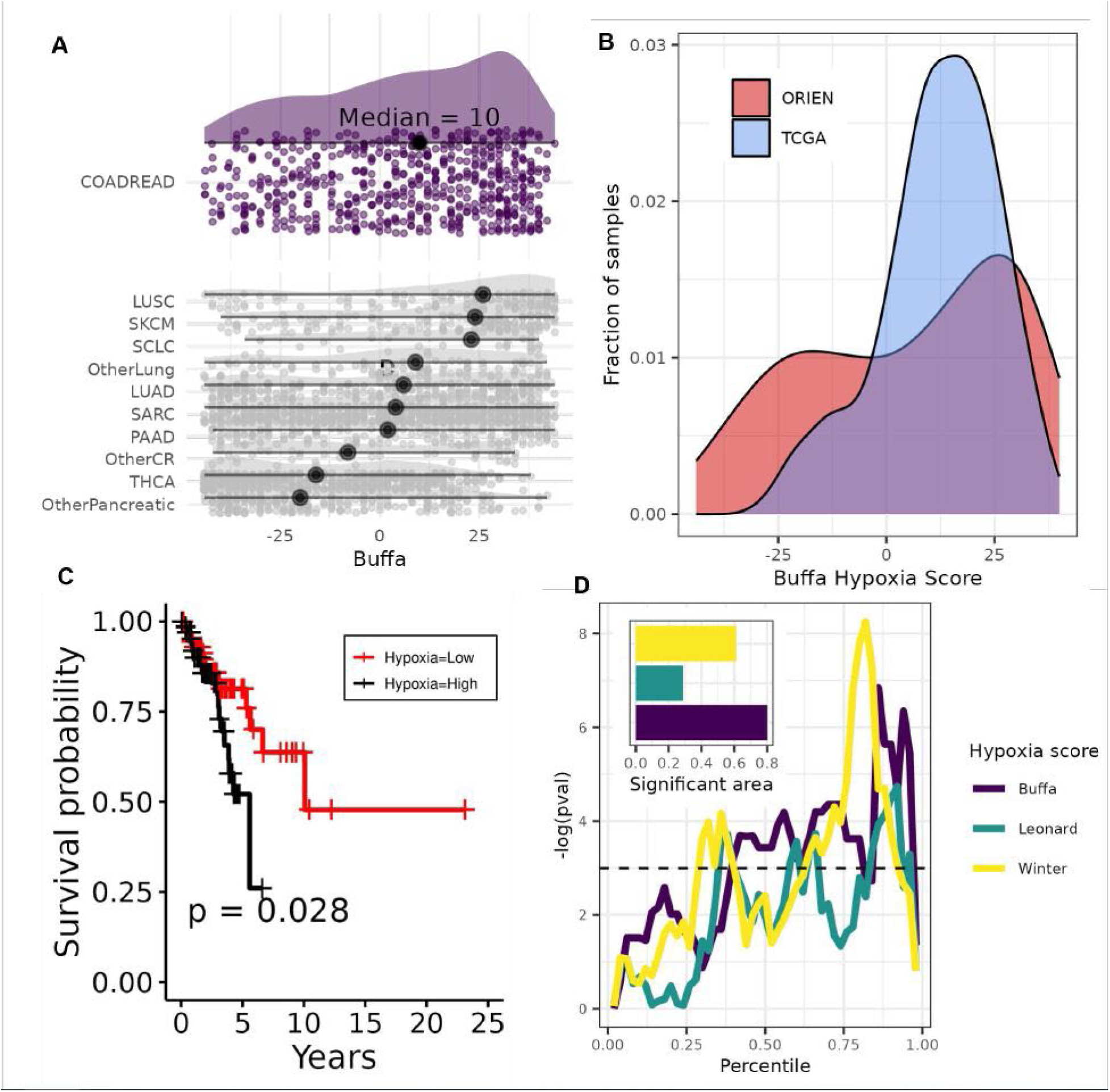
Hypoxia exists in CRC and predicts for poor patient outcome after XRT. **(A)** Hypoxia expression signature scores for the ORIEN dataset across tumor sites showing a wide distribution for COAD and READ samples. The median values are indicated with a black circle. **(B)** Comparison of Buffa score histogram of ORIEN with TCGA CRC datasets showing a very similar mode in both, but with a wider distribution for ORIEN. **(C)** For ORIEN patients receiving radiotherapy, stratification of patient population into upper and lower hypoxia tertiles shows that hypoxic tumors have worse patient outcomes. **(D)** Hoyd plot of hypoxic signatures shows clinical significance of hypoxia scores over a range of stratifications.

We downloaded RNA-seq data from 141 CRC patients available through the ORIEN network and calculated hypoxia scores (23) for each sample. Colorectal adenocarcinoma (COADREAD) ranks among the more hypoxic cancer types with a median hypoxia score of 12. The distribution of hypoxia scores ranges from lowest in thyroid cancer (median hypoxia expression score of 16) to highest in lung squamous cell cancer (median hypoxia expression score of 26). We then cross-validated the ORIEN COADREAD hypoxia scores by comparing them with the publicly available TCGA COADREAD dataset. We found substantial overlap in the distribution of scores between the independent patient datasets, confirming the robustness of this analysis and indicating that the many COADREAD samples were distributed within the high-end hypoxia expression score (23) region (**Figure 1B**). Current molecular models of radiation-induced DNA damage specify that oxygen is needed for the “fixation” of damage; thus, hypoxia directly impairs the efficiency of radiation therapy (27). Therefore, we tested whether elevated hypoxia scores would correlate with poor survival of CRC patients treated with radiation therapy. ORIEN patients who underwent radiation therapy were stratified at the median hypoxia expression score (23), and KM analysis shows that a high hypoxic score significantly correlates with poor overall patient survival (p=0.028) (**Figure 1C**). For a more robust survival analysis, we evaluated whether stratification strategy influences significance within these COADREAD patients. To this point, we stratified this dataset of hypoxia expression score (23) tumors over a range of different values and analyzed the impact on the significance of outcome prediction. **Figure 1D** shows this analysis for 3 published hypoxia gene scores. The predictive ability of a high hypoxia expression score (23) is significant for survival over a wide range of stratification points, and the hypoxia expression score (23) appears to be the most discriminating of the signatures.

Significant evidence suggests that the composition of the gut microbiome correlates with the initiation and progression of CRC and affects tumor sensitivity to anti-cancer therapy (15,28). Accordingly, we evaluated tumor microbiota composition in ORIEN and TCGA patient samples using the exogenous sequences in tumors and immune <u>c</u>ells {exotic} tool for quantifying microbe abundances in tumor RNA-seq data (24). To identify taxa associated with high levels of tumor hypoxia, we stratified patient samples into tertiles of increasing hypoxia scores (23) and quantified log2 abundance fold change between the bottom- and top-ranked tertiles. This analysis was performed on both ORIEN and TCGA CRC datasets, and only those microbes that were significant in both datasets were identified as hypoxia-trophic or -phobic. A volcano plot of significant strains is shown in **Figure 2A**, and a bar graph with the magnitude of the effect is shown in **Figure 2B**. Results for all log2 fold changes are available in **Table S1**. Interestingly, *Fusobacterium nucleatum* has been associated with poor outcomes in patients with CRC (15,29) and is elevated in hypoxic tumors.

**Figure 2.**
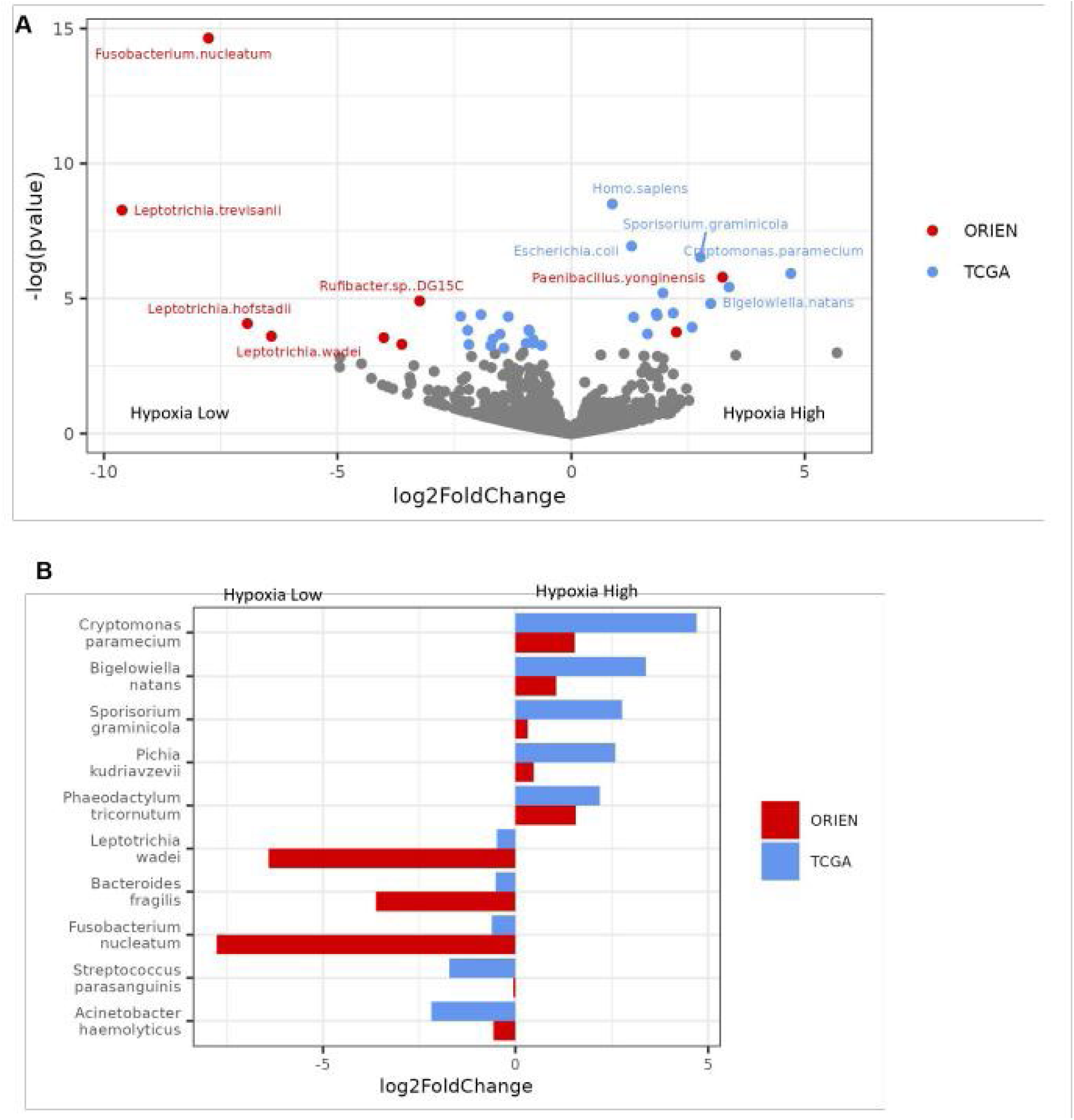
Hypoxia score associated with significant differences in microbial populations in CRC. **(A)** Volcano plot of microbes from ORIEN and TCGA tumors showing strain differences when tumors are stratified by high vs. low hypoxia scores (tertiles). **(B)** Fold change of microbes that show hypoxic enrichment in both ORIEN and TCGA datasets.

We next analyzed the intratumoral microbial populations in these patients for strains that correlated with the outcome of radiotherapy for CRC. We identified 11 microbes whose presence correlated with significantly worse outcomes after radiotherapy **(Figure 3A)**. Complete results for the survival analyses are available in **Table S2**. Strains showing the strongest statistical correlation included *Candida glabrata*, a fungus, and *Fusobacterium canifelinum* and *Bulleidia* sp zg 1006, both bacteria. *Fusobcterium canifelinum* is an anaerobic, gram-negative bacilli that is not generally present in a healthy human gut microbiome but typically occurs in the oral cavity (30). The closely related *Fusobacterium nucleatum* has been identified in recurrent CRC, and the presence of the *Fusobacterium* strain promotes CRC development (31,32) and chemoresistance (16). We therefore asked whether the combined presence of hypoxia and *F. canifelinum* affected treatment outcomes in CRC patients undergoing radiation therapy. Analysis of CRC patients treated with radiation therapy identified that the presence of *F. canifelinum* or a high hypoxia score (23) is significantly associated with reduced overall survival (p-values : *F. canifelinum* = 0.005, *F. canifelinum* interacting with hypoxia < 0.001) (**Figure 3B**).

**Figure 3.**
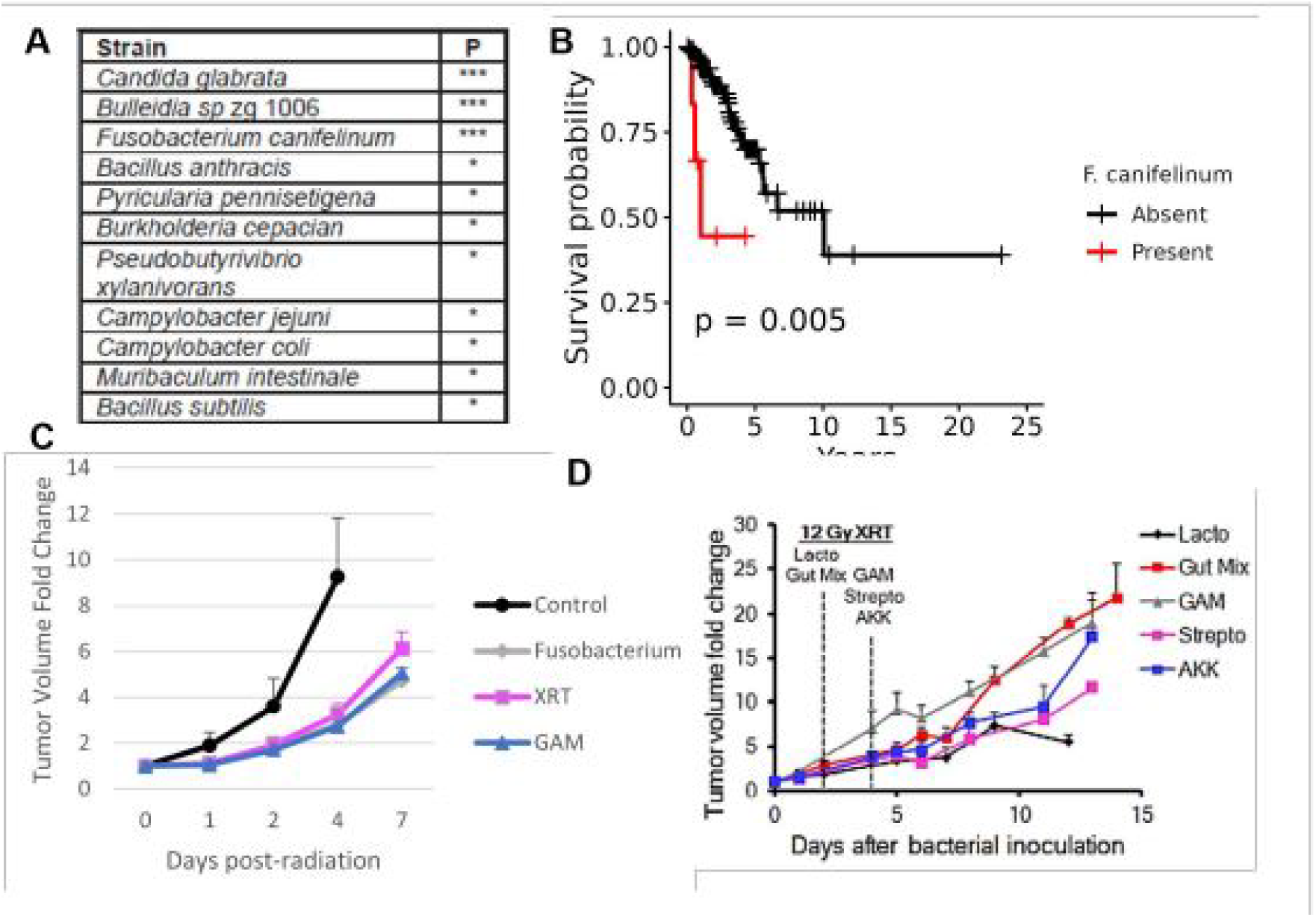
Presence of certain microbes in ORIEN tumors correlate with worse outcomes after radiotherapy. **(A)** List of strains whose interactions with hypoxia in the tumor are significantly associated with survival outcomes in ORIEN patients who received radiation treatment. **(B)** Kaplan-Meier survival curve showing association between the presence of *F. canifelium* and patient outcome. **(C)** Model CT26 tumors grown in immune-deficient mice and inoculated with *Fusobacterium* shows no increase in tumor growth delay after radiotherapy. **(D)** Similar model tumors inoculated with several other bacterial strains showing significant increase in tumor growth delay after radiotherapy.

To experimentally validate our bioinformatic observations in vivo, athymic nude mice were inoculated heterotopically with the mouse CRC line MC38. Upon reaching 100 mm^3^, which is the volume typically associated with the formation of a hypoxic core, MC38 tumor-bearing mice were randomized into groups receiving either nothing as control, intratumoral bacterial growth medium (GAM), or cultured *F. nucleatum* grown in GAM to achieve a ratio of 1:1 bacteria to tumor cells. After 48 hours, mice were either sham irradiated or given one dose of 8 Gy external beam radiation using the SARRP. **Figure 3C** shows no significant change in tumor response to therapy with the introduction of *F. nucleatum*.

We therefore decided to test other bacterial strains for effects on radiation response and repeated this experiment using several common gut bacteria strains: 1) microbial coculture representative of a healthy intestinal microbiome (gut mix); 2) *Lactobacillus spp*.; 3) *Streptococcus spp*.; or 4) *Akkermansia spp*., also by intratumoral injection at a 1:1 ratio of microbe to tumor cell, followed 48-72 hours later by a 12 Gy dose of local radiation. **Figure 3D** shows that injection of *Lactobacillus* or *Streptococcus* bacterial strains into heterotopic CRC tumors resulted in a pronounced increase in tumor growth delay after radiation. Taken together, these findings support the idea that intratumoral microbes can influence the response of model tumors to radiotherapy in a strain-dependent manner.

We next investigated the impact of a hypoxic TME on spontaneous tumor microbial colonization in model tumors in mice. We inoculated a cohort of 10 BALB/c and 10 athymic nude host mice with the CT26 murine CRC cell line. The mice received CT26 cells subcutaneously in the flank, and their tumors were harvested upon reaching 500 mm^3^ for nucleic acid extraction. Hypoxia scores (23) were calculated from tumor RNA for each sample, and the tumors were stratified at the mean hypoxia score (23) into normoxic (low hypoxia expression score) and hypoxic (high hypoxia expression score) tumors in each mouse host (**Figure 4A**). We used metatranscriptomic sequencing of exogenous RNA to identify microbial strains enriched in hypoxic tumors. **Figure 4B** shows the bacterial burden of the tumors by calculating the fraction of RNA reads that were bacterial. In the immune-competent model, the bacterial burden was equivalent in both low- and high-hypoxia tumors. However, in the tumors grown in nude mice, the burden was higher in hypoxic tumors, potentially due to the predominance of anaerobic strains (**Figure 4C**). A comparison of the bacterial phyla identified in each group shows similarities in the tumors and the prominence of *Firmicutes* phylum (**Figure 4C**). Microbes were identified that were significantly enriched in the normoxic and hypoxic tumors for each host (**Figure 4D**). **Table S3** contains the results summarized in **Figure 4D**. Interestingly, there was an overlap of 3 of the strains found preferentially in the hypoxic tumors in both humans and mice (**Figure 4E**). The analyses of these microbes in each dataset are presented in **Table S4**.

**Figure 4.**
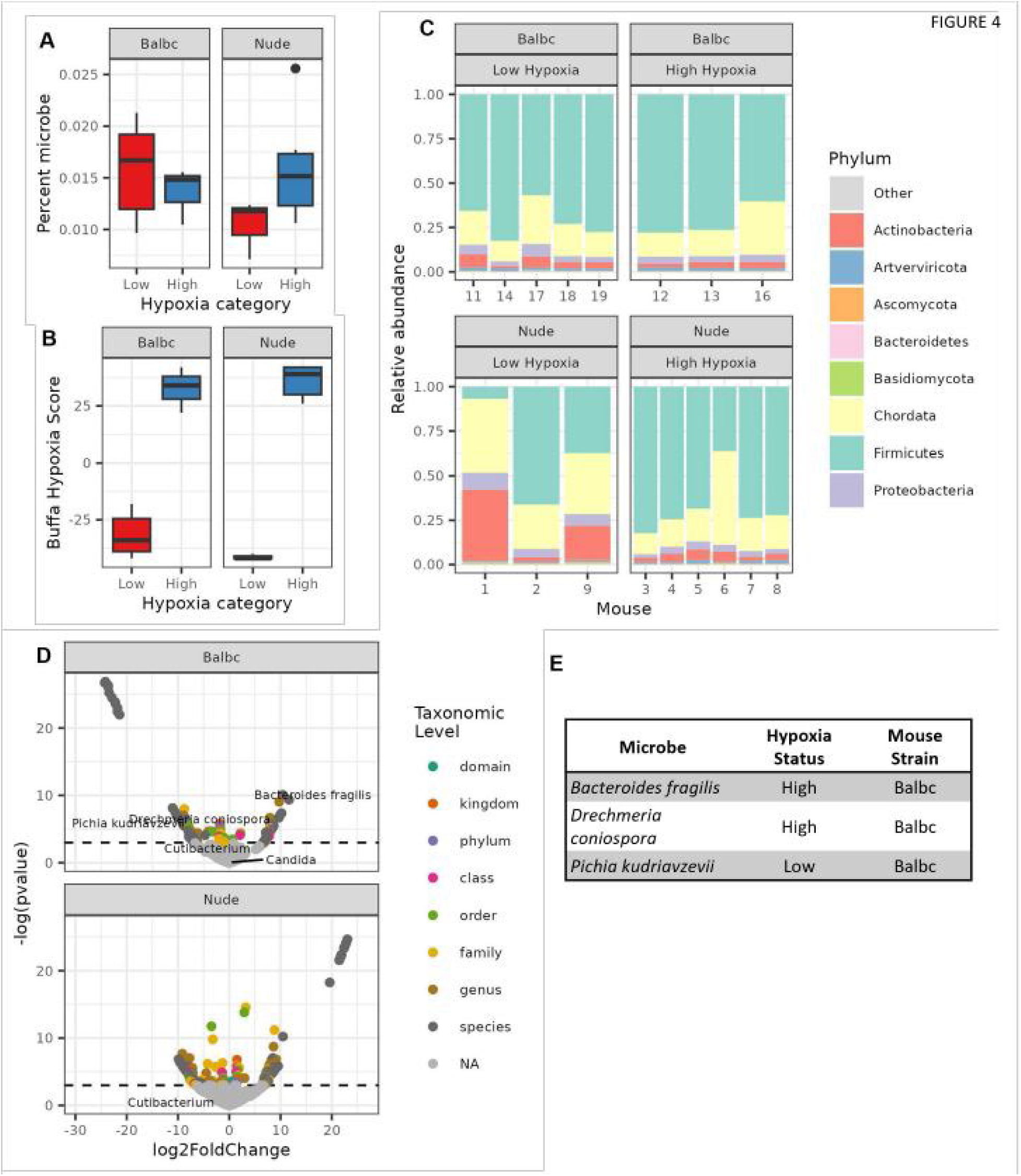
Model tumors also have oxygen-dependent microbial strain differences. **(A)** CT26 CRC tumors grown in immune deficient hosts have increased microbial burden with increased hypoxia. **(B)** CT26 CRC tumors grown in immune deficient or immune competent hosts can be stratified into low and high hypoxia groups by tumor Buffa score. **(C)** Metatranscriptomic identification of microbial strains spontaneously colonizing individual tumors. **(D)** Volcano plot of strains identified in C with tropism for tumor of either high or low hypoxia score. **(E)** Microbes found to be enriched in both mouse and human hosts with hypoxia status.

However, the most consistent colonization of all tumors was observed with the genus *Cutibacterium* from the phylum *Actinobacteria* (**Figure 4D**). *Cutibacterium* is gram-positive anaerobic bacilli typically found in the skin and associated with acne vulgaris; however, emerging studies identified a possible role for the microbe in prostate cancer immunosuppression (33). Based on the prevalence of *Cutibacterium* in all analyzed samples, we performed transcriptomic analysis of this species in both the BALB/c and nude mouse hosts. We analyzed how the same microbial strain responds to these TME variables (tumor oxygenation as determined by hypoxia expression score) in different mouse hosts. Differential microbial RNA analysis revealed significant differences between microbial gene expression in normoxic versus hypoxic tumors, indicative of microbial adaptation to TME changes (**Figures 5A and 5B, Table S5**). Of the 1,295 identified RNA species, we found 48 to be significantly different in the normoxic versus hypoxic tumors in both datasets (39 induced by hypoxia and 9 repressed). The 34 with Uniref90 annotations are listed in **Figure 5B** in order of significance.

**Figure 5.**
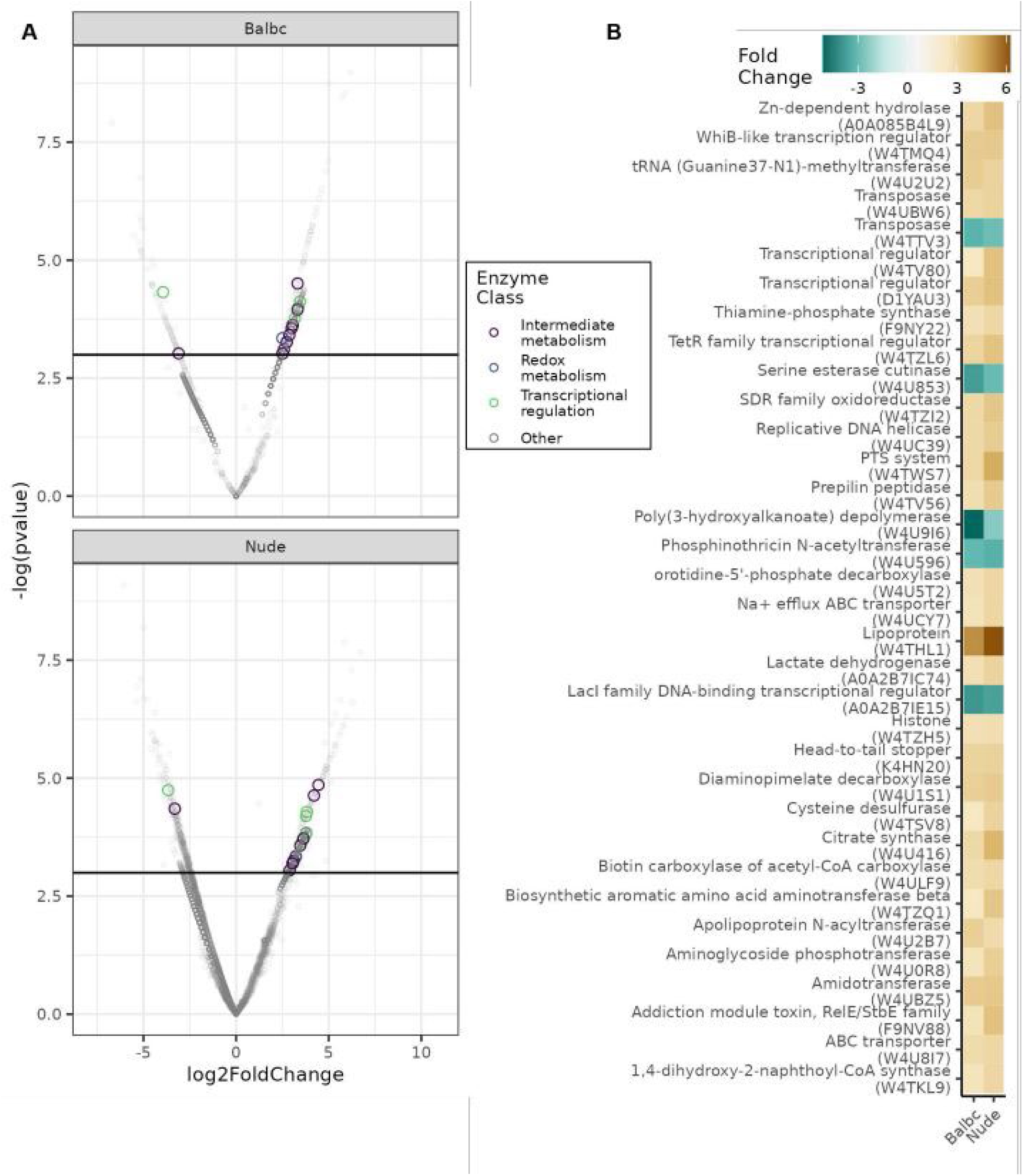
Oxygen-responsive gene expression of *Cutibacterium* within model tumors. **(A)** Volcano plot of metatranscriptomic analysis of transcripts from *Cutibacterium* found in low or high hypoxia score CT26 tumors grown in either immune-deficient or immune-competent host animals (1,295 total transcripts identified). **(B)** Table of the 34 annotated transcripts that showed significant concordant changes to hypoxia score in both datasets (49 total: 40 positive, 9 negative).

Even with the paucity of annotation in this strain, we find that adaptive changes included genes primarily involved in metabolism but also several involved in oxygen/redox stress, DNA metabolism, and several transcriptional regulators. Hypoxic induction of lactate dehydrogenase or citrate synthase may be analogous to what is seen in higher species adapting their metabolism to reduced oxygen environments. One of the more interesting induced transcriptional regulators is the WhiB family member, which contains a redox-sensitive [4Fe-4S] cluster. The WhiB-like (Wbl) protein family is exclusively found in *Actinobacteria* and has been reported to be nitric oxide and oxygen-responsive in other strains (34). This transcriptional regulator may be analogous to the hypoxia-responsive HIF1 transcriptional regulator found in higher species that is largely responsible for transcriptional adaptation to hypoxia. The sum of these findings indicates that intratumoral oxygen tension can both favor the growth of specific microbial populations and elicit adaptive transcriptional responses in microbes that can grow in both normoxic and hypoxic tumors.

## DISCUSSION

Several lines of evidence show that host-microbiome interactions can affect the response to anti-cancer therapy (14,35–37). The balance between bacterial and fungal microbes and the metabolites produced by these microbes can affect response to radiation therapy (38,39). Chronic intestinal inflammation associated with microbial dysbiosis and an intrinsically hypoxic microenvironment creates a niche permissive to microbial colonization of all ranges of partial oxygen pressure (pO_2_) within the tumor. In line with this expectation, we observed that tumor microbiome composition shows distinct patterns as a function of tumor oxygen levels in CRC patient datasets.

Under normal conditions, the intestinal microbiome consists of 1,000+ species of bacteria, most of which are obligate anaerobes (40). Intestinal dysbiosis triggers the rapid expansion of conditional anaerobic pathogens that can disrupt the intestinal mechanical barrier and result in luminal microbes colonizing the tumor. Mean pO_2_ levels within the GI tract range from 42-71 mmHg (7-10% O_2_) in the colonic muscle wall to 3 mmHg (0.4% O_2_) in the sigmoid colon, similar to 2.5 mmHg levels typically observed in CRC tumors (41,42). Distribution characteristics of conditional pathogens in both mouse hosts show an overrepresentation of the *Firmicutes* phylum, similar to observations made in human CRC (43).

Much investigation has gone into the composition of the intratumoral immune system to establish its role in determining which microbes can colonize a tumor (44). However, there are fewer studies of the metabolic parameters that influence what microbes find a tumor hospitable. The partial pressure of oxygen has been long recognized as an important component of culture media for different microorganisms. Bacteria have been found that range from obligate anaerobes that can only grow in the absence of oxygen to aerobes that grow well in an atmosphere with 21% oxygen. Often, this categorization is due to the presence or absence of enzymes within the microbial genome that are capable of metabolizing toxic oxygen species, such as superoxide dismutase or catalase, which can facilitate growth in an oxidizing environment. Many microbes fall between these extremes and can be categorized as facultative anaerobes or microaerophiles capable of growth in low- or no-oxygen environments. These microbes may be most well adapted for growth in intermediate to low oxygen, which is often found within the TME.

Similarly, while much attention has been given to what strains of bacteria exist within the tumor, relatively less study has been completed on the physiologic state of intratumoral microbes. Microbes could have a very different impact on the tumor due to the expression or silencing of different genes. Our data suggest that differing levels of microbially produced lactate or citrate may be present in the hypoxic tumors relative to the more well oxygenated tumors. Such metabolites may interact with and change the behavior or morphology of the tumor. For example, tumor lactate has been shown to impact macrophage polarization (45). Such signals from microbes have also been shown to impact tumor cell response to radiotherapy (39). In our study, we observed several microbial strains correlated with worse outcomes from radiation therapy in rectal cancer. Notably, *C. glabrata* and *F. canifelinum* were identified, both of which share close taxonomic similarity to microbes previously related to cancer: *Candida albicans* and *Fusobacterium nucleatum,* respectively. *C. albicans* has been linked to oral squamous cell carcinoma (46) and radioresistant rectal cancer (47) and was shown in a large study to be predictive of metastatic disease across multiple GI tumors (48). Additionally, previous studies have reported that species from the *Fusobacterium* genus are associated with elevated inflammatory response in the colorectal mucosa (49–51). This raises the question of whether the host’s immune response to microbial dysbiosis affects radiation therapy outcomes.

Experimentally, we observed that immunocompromised BALB/c and nude mice hosts show distinct acute immune responses following intratumoral inoculation with several microbial strains found in human CRC (52). The inoculated mice exhibited mild symptoms of acute immune response associated with changes in tumor growth and response to radiation therapy. We identified that the presence of common commensal bacteria *Lactobacillus acidophilus, Akkermansia municiphila,* and *Streptococcus thermophilus* in the tumor enhances the therapeutic efficacy of local radiation therapy while inoculation with *Fusobacterium nucleatum* or microbial coculture, representative of a healthy intestinal microbiome, does not potentiate radiation therapy. These observations support the model that the presence of *Lactobacillus*, *Akkermansia,* or *Streptococcus* genera might be predictive of better outcomes of radiation therapy in CRC patients, while the presence of *Fusobacterium* predicts worse outcomes (*F. nucleatum* in previous publications, and *F. canifelinum* in our study).

Perhaps the most intriguing findings come from the metatranscriptomics analysis of conditional pathogen enrichment in hypoxic versus normoxic CT26 CRC tumors in BALB/c and nude mice. We identified that *Cutibacterium*, a typically anaerobic opportunistic pathogen, is most consistently colonizing both the normoxic and hypoxic tumors, displaying wide adaptive flexibility toward different pO_2_ environments. Transcriptomic analysis revealed a systemic reprogramming underlying the adaptive response of the pathogen to varying pO_2_ levels. These results support the model that opportunistic pathogens colonizing CRC tumors can respond to environmental oxygen levels with adaptive gene expression changes, providing a strong rationale for investigating microbial metabolism in future studies. These results also indicate that it may not be just the presence or absence of a specific microbe within a tumor that dictates its effect on clinical outcome but the physiological state of the microbe that can be influenced by the microenvironment in which it resides.

## SUBMISSION SECTION

Precision Medicine & Biomarkers

## CODE AVAILABILITY

The code to reproduce all analyses and figures is available at https://github.com/spakowiczlab/recrad

## AUTHOR CONTRIBUTIONS

Conceptualization: DS, ND

Resources: DS, LAR, CC, YZ, RDD, CMU, SH, MC, AAT, EAS, API, MM, AEGO, ACT, QM, GR, BS, JZ

Data curation: RH, CEW, MB, MK, JZ, FC

Software: RH, CEW, DS

Formal analysis: RH, CEW, DS, MB, JZ, FC

Supervision: CEW, JZ

Validation: RH

Investigation: RH, MB, MK, JZ, FC, KB, MK

Visualization: RH, CEW, MB, MK

Methodology: RH, CEW, DS

Project administration: DS, ND

Writing - original draft: CEW, DS, MC, ND, MB

Writing - review and editing: RH, CEW, DS, LAR, CC, YZ, RDD, CMU, SH, MC, AAT, EAS, API, MM, ND, GT, MH, NJ, MB, MK, DJG, IE, ACT, QM, GR, BS, JZ, IP, FC, KB, MK

## ACKNOWLEDGMENTS

This work was supported by the Pelotonia Institute of Immuno-Oncology (PIIO). The content is solely the responsibility of the authors and does not necessarily represent the official views of the PIIO. The authors acknowledge the support and resources of the Ohio Supercomputer Center (PAS1695, PCON0005). We would like to thank Angela Dahlberg, Editor, Division of Medical Oncology at The Ohio State University Comprehensive Cancer Center, for editing and proofreading the manuscript.

